# Development of a Pharmacokinetic (PK) Mouse Serum GLP ELISA for an Anti-CD19–Anti-CD3 Diabody

**DOI:** 10.1101/2025.03.19.644217

**Authors:** Tonny Johnson, Armin Ahnoud, Elisabeth DiNello, Peter Ralph, Timothy P. Cripe, Pin-Yi Wang, Tania L. Weiss

## Abstract

GP101 is a single chain diabody composed of Fv regions of antibodies directed to CD3 and CD19. It is designed to mimic commercial blinatumomab (Blincyto®) that simultaneously co-engages patient T lymphocytes and CD19 positive B cell leukemias and lymphomas, facilitating their targeted destruction. This study details the purification of GP101 protein and development of a highly sensitive ELISA for its detection, achieving a sensitivity of 15pg/mL in 25% mouse serum. This ELISA has been validated according to GLP standards.

## 1. Introduction

A sensitive ELISA method is crucial for drug development in clinical trials. It measures drug concentration in biological samples such as blood and plasma over time, which is essential for understanding drug absorption, distribution, metabolism, and excretion. This information helps determine optimal drug dosing and scheduling (1,2).

An adeno-associated virus (AAV)-based single-dose immunogene therapy has been developed to address the current limitations of CAR-T cell and bispecific antibody-based T-cell immunotherapy (3) for B cell leukemias and lymphomas. The AAV vector, VNX-101, expresses a gene encoding the gene product (GP)101, a blinatumomab-mimic (αCD19-αCD3 diabody). This vector offers advantages over protein therapies that require continuous administration, and CAR-T cell therapies that require individualized patient manufacturing and pose risks such as cytokine storms and other unpredictable side effects (3). This technology has been licensed by Vironexis Biotherapeutics, Inc. to develop further, which is currently being tested in a phase 1 clinical trial (NCT06533579). A previous report describes an ELISA for Blincyto with a sensitivity of 50 -100 ng/mL (4), which is inadequate for detecting effective GP101 drug concentrations in preclinical or clinical trials as its serum concentrations at FDA recommended doses are in the 0.5 – 3.0ng/ml range.

The challenge in developing a highly sensitive ELISA for blinatumomab may be due to its dissociation from its target proteins, CD3 and CD19. The CD3 has approximately 183-fold lower affinity for Blincyto compared to CD19, Kd of 2.6 × 10^−7^ M vs 1.49 × 10^−9^ M, respectively (5).

This article describes the purification of recombinant GP101 protein and the establishment of a highly sensitive ELISA for its quantification. A standard GP101 was produced by transfection of a plasmid encoding GP101 into Expi293 cells, and by a single-step purification by affinity chromatography. The ELISA had a detection limit of 15pg/mL (0.3 femtomoles/mL) in 25% mouse serum. The method was validated for GLP PK studies in VNX-101 treated mice.

## 2. Materials and Methods

### 2.1. Materials

A plasmid containing the gene for GP101, essentially pAAV[Exp]-CAG > {sec2A-CD3xCD19, is described by Cripe et al. (3). The following reagents were purchased, Expi293 cells (Thermo Fisher), Protein Concentrators spin column, 10K MWCO (Pierce cat# 88513), protein CD-19 conjugated magnetic beads (ACROBiosystems), NuPAGE™ 4 to 12%, Bis-Tris, Mini-Protein Gel (ThermoFisher), CD3 protein (ACROBiosystems) and CD19 protein conjugated to human Fc (R&D Systems), HRP-anti HumanFc (Abcam), QuantaRed (ThermoFisher), normal mouse serum (Sigma), and Blincyto (Amgen).

### 2.2. Production of recombinant anti-CD19–anti-CD3 diabody (GP101)

The plasmid containing the gene for an GP101was used to transfect Expi293 cells according to the manufacture’s recommendations. Cell culture supernatants were collected daily for six days for analysis.

### 2.3. Blinatumomab ELISA

The blinatumomab ELISA method of Moore *et al*. (4) was followed. Aliquots of cell culture supernatants were removed from flask each day for six days and were screened for the expression of GP101.

### 2.4. Purification of recombinant GP101

Three-day cell culture supernatants were concentrated with a spin column. The GP101 was purified using CD19-coupled magnetic beads according to the manufacture’s protocol. The concentration of purified GP101 was determined by measuring the optical density (OD) at 280nm, using an extinction coefficient of 2.26 at 1mg/mL, calculated based on its amino acid composition. The molecular weight and purity of the purified GP101 was estimated by SDS-PAGE electrophoresis. This material is cited as GP101 Reference Standard and is stored in aliquots at -70° C.

### 2.5. Development of the highly sensitive GP101 ELISA

CD3εδ protein (ACROBiosystems) was coated onto wells of a black ELISA plate (Thermo Fisher) and incubated overnight at 4°C. The wells were blocked with 1% casein in PBS (Thermo Fisher). The GP101 samples were pre-incubated with CD19-human Fc (R&D Systems) for 1 hour at 37°C in the presence of 2% casein as an assay diluent. The GP101/CD19 mixture was added to the blocked wells and incubated for an additional hour at 37°C.

GP101 standards were prepared in 2% casein-Tris for buffer-only ELISA experiments. The ELISA for mouse serum, the GP101 standard was diluted in assay diluent containing a final concentration of 25% pooled normal mouse serum (5). After washing the wells, HRP-conjugated anti-human IgG-Fc was added, followed by QuantaRed™ substrate, which was incubated according to the manufacturer’s instructions, with modifications as necessary.

Fluorescence was measured using a Cytation 5 plate reader (Agilent). All conditions were tested in triplicate. Relative Fluorescence Unit (RFU) were analyzed using a four-parameter (4P) curve fit in Gen 5 software to generate the standard curve and determine GP101 concentrations (pg/mL) in test samples.

## 3. Results

### 3.1. Expression screening of GP101 in transfected Expi293 cells

The plasmid containing the gene for GP101 was transfected into Expi293 cells and supernatants were tested on various days for GP101 concentration using the blinatumomab ELISA. The relative luminescence based upon the GP101 concentration, show that GP101 expression reaches its peak on the third day, after which no further increase is observed (Fig. 1).

**Fig. 1:**
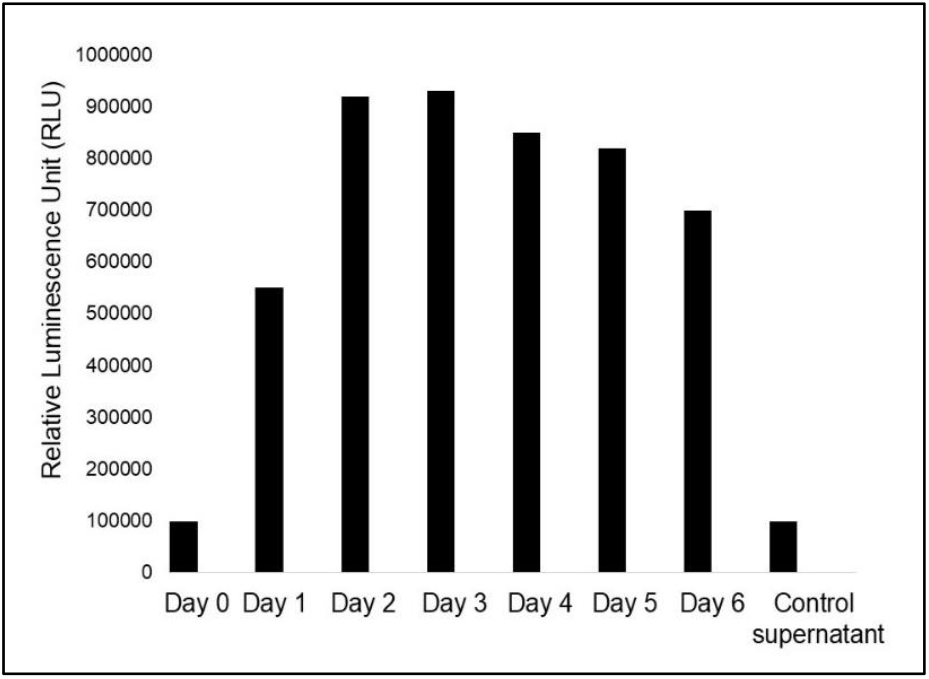
Expression screening of GP101 in Expi293 cells.

### 3.2. Single-step affinity chromatography purification of GP101 to near homogeneity

The cell supernatant collected on the third day of culture was used for GP101 purification using CD19-coupled magnetic beads. The purification process was monitored using SDS polyacrylamide gel electrophoresis. The results indicate that the majority of the eluted GP101 was collected in the first two fractions, with a major band of approximate molecular weight (MW) ∼55kDa at over 90% purity (Fig.2A). When the gel is overloaded, a minor band at approximately ∼130 kDa is seen comprising approximately 3% of the total protein as seen by Coomassie staining and scanning using ImageQuant for quantitation.(Fig. 2B).

**Fig. 2A.**
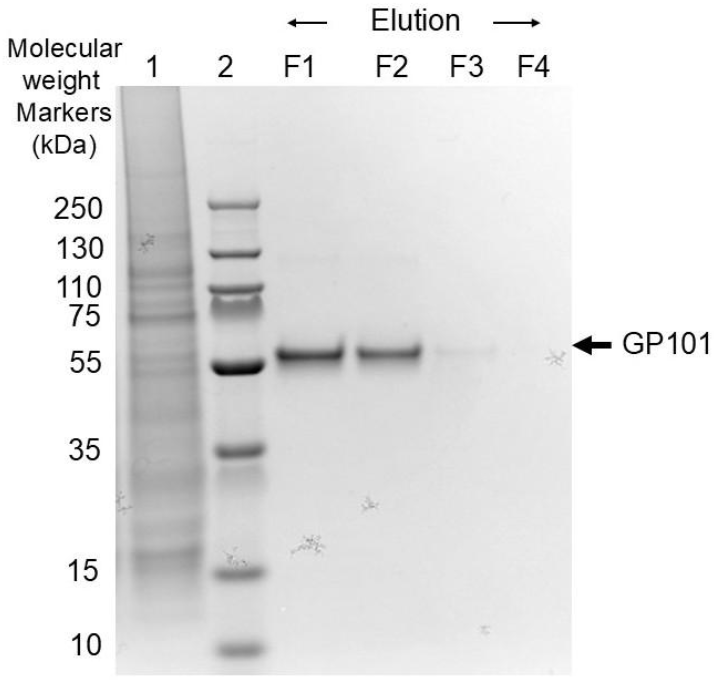
SDS-PAGE analysis of GP101 purification. **Lanes: 1**. Cell culture supernatant expressing GP101. **2**. Molecular weight (MW) markers. **F1**. Eluted fraction #1. **F2**. Eluted fraction #42. **F3**. Eluted fraction #3. **F4**. Eluted fraction #4.

**Fig. 2B.**
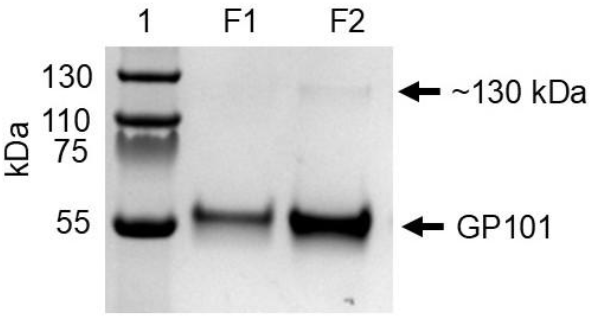
ImageQuant analysis of purified GP101 fractions. **Lanes: 1**. Molecular weight (MW) markers. **F1**. Eluted fraction #1. **F2**. Eluted fraction #2.

**Fig. 2C.**
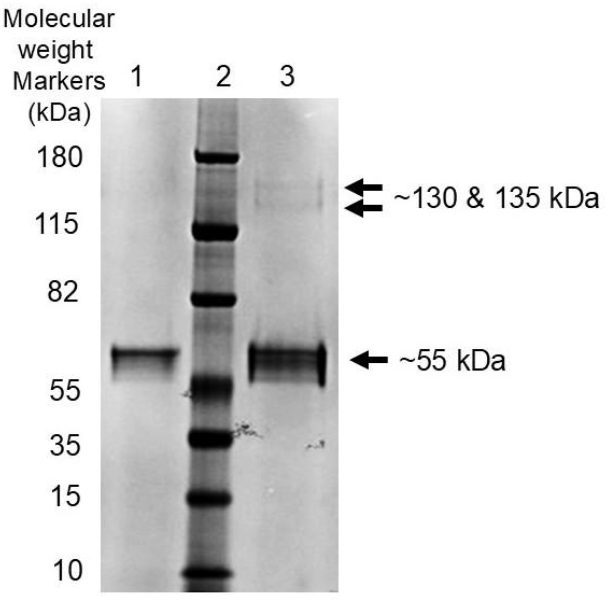
SDS-PAGE analysis of purified GP101 and Blincyto®. **Lanes: 1**. Purified GP101. **2**. Molecular weight (MW) markers. **3**. Blincyto®.

A comparison of GP101 and Blinycto by SDS-PAGE and ImageQuant analysis shows that GP101 molecular weight is similar to Blincyto (∼54 kDa). Blincyto shows two faint bands at approximately 130 to 135 kDa comprising approximately 3% of the total protein (Fig. 2C); GP101 was not overloaded so the high MW band is not seen in this gel.

Purification of GP101 was carried out through three independent experiments. The concentration, identity, and purity of the purified protein were assessed in each experiment using absorbance at 280nm, SDS-PAGE, and a GP101 specific ELISA. We successfully purified approximately 400µg of GP101 protein with >90% purity, as confirmed by ImageQuant analysis.

### 3.3 Development of an ELISA method for the detection and quantitation of GP101

A series of experiments were performed to evaluate various components and conditions for ELISA development. Different blocking reagents including gamma globulin-free bovine albumin, casein, and Fish Serum Blocking Buffer (ThermoFisher) were studied, ultimately finding casein to be the most effective blocker (data not shown). For coating and detection of proteins, we tested both human CD3 and CD19 proteins, available in various constructs. The CD3 (CD3εδγ) performed better as a coating protein than CD19, based on background signal and positive signal strength. We selected a CD19-human Fc construct for detection, paired with an HRP-conjugated anti-human Fc secondary antibody. To optimize signal generation, we evaluated multiple enzyme-substrate combinations, including various HRP substrates (TMB, Super Signal Pico PLUS Chemiluminescent Substrate, Amplex Ultra Red, and QuantaRed™) as well as alternative systems such as alkaline phosphatase and biotin-streptavidin HRP. The final method is illustrated in Fig. 3.

**Fig. 3:**
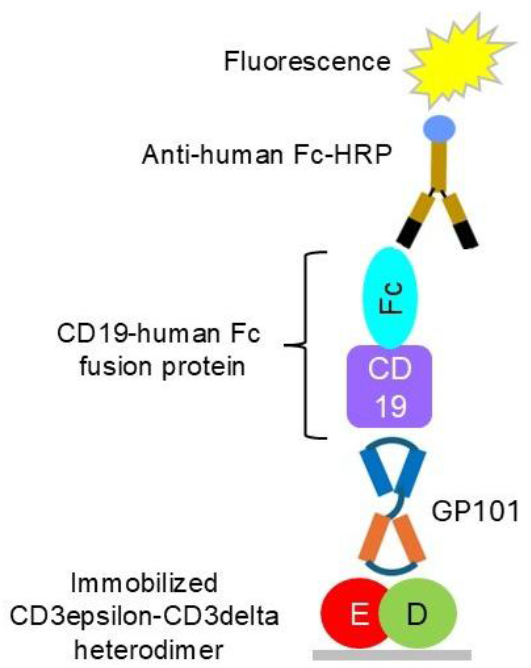
Schematic diagram of the GP101 ELISA.

In brief, GP101 is pre-incubated with CD19-Fc to form a GP101:CD19-Fc complex. This complex is then captured by a CD3 heterodimer protein immobilized on a 96-well plate, with unoccupied sites blocked by casein. Detection is performed through binding of anti-Fc-HRP. The HRP catalyzes QuantaRed™ substrate producing a fluorescent signal.

For quantitation of GP101 in mouse serum, we conducted tests using varying concentrations of mouse serum to determine the optimal conditions for accurate measurement. Among the different concentrations tested, 25% mouse serum demonstrated superior performance in terms of sensitivity, as indicated by higher signal-to-noise (S/N) ratio, accuracy and precision. Higher concentrations of mouse serum were not suitable (data not shown). The sensitivity of the ELISA is 15pg/mL (0.3 femtomoles/mL) GP101 in 25% mouse serum (Table 1). The upper limit is >>500pg/mL.

**Table 1.** Sensitivity of GP101 ELISA in 25% mouse serum: 15pg/mL is chosen as the sensitivity limit based on S/N and precision of 3 replicates (%CV).

| GP101<br>(pg/mL) | 500 | 250 | 125 | 50 | 30 | 15 | 0 |
| --- | --- | --- | --- | --- | --- | --- | --- |
| Average Relative<br>Fluorescence unit<br>(RFU) | 33614 | 18937 | 8856 | 5905 | 3965 | 3237 | 833 |
| STDEV | 1447 | 1074 | 187 | 278 | 423 | 88 | 65 |
| %CV | 4 | 6 | 2 | 5 | 11 | 3 | 8 |
| S/N | 40 | 23 | 11 | 7 | 5 | 4 | 1 |

### 3.4 Development of a validated GP101 ELISA method for GLP pharmacokinetic (PK) studies in mice

The US FDA guidelines for ELISA specify five quality controls: Upper and Lower Limits of Quantitation (ULOQ, LLOQ), and three internal controls during validation. For robustness of the assay, we picked 40pg/mL as the LLOQ and 10,000pg/mL as the ULOQ. The lower, mid and high internal QC controls were chosen to be 100, 640, 4,000pg/mL in 25% mouse serum. The accuracy of these controls across three independent ELISA experiments was 84% or higher. The precision of replicates varied from 1%CV to 13%CV (Table 2).

**Table 2.** Accuracy and precision of the 3 independent ELISA experiments at ULOQ (10,000pg/mL), LLOQ (40pgL), and three QC samples (4,000, 640 and 100pg/mL).

|  | GP101 concentrations (pg/mL) |  |  |  |  |
| --- | --- | --- | --- | --- | --- |
|  | 10000 | 4000 | 640 | 100 | 40 |
|  | Accuracy |  |  |  |  |
| Mean | 84% | 92% | 92% | 85% | 89% |
|  | Precision of replicates |  |  |  |  |
| Mean CV | 3% | 13% | 1% | 10% | 9% |

An example of the purified GP101 standard 4P curve fit is shown in Figure 4.

**Figure 4.**
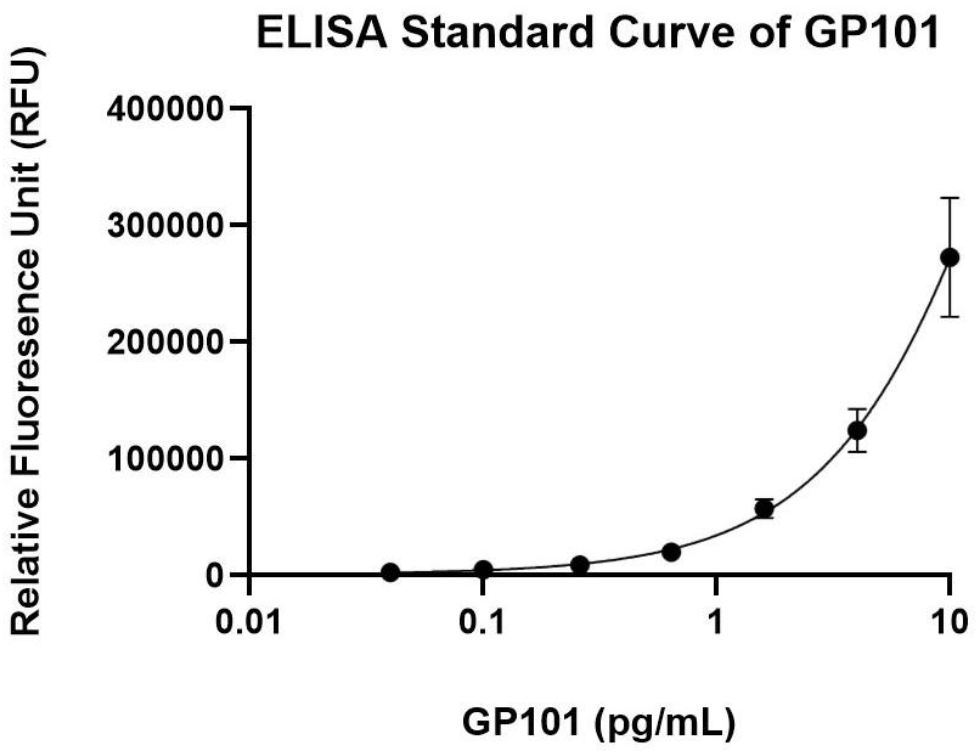
Highly sensitive ELISA standard curve for validation.

Additionally, to assess background response, 10 individual mouse sera from untreated animals were tested as well as a pooled normal mouse serum without GP101. These negative control signals were consistently below the lower limit of quantitation (LLOQ). The pooled serum negative control is included in every PK assay.

## 4. Discussion and Conclusion

Our affinity-purified GP101 protein showed >90% purity based on SDS-PAGE and ImageQuant analysis, confirming the efficiency of our single-step purification strategy using CD19-coupled magnetic beads. The observed molecular weight of approximately (∼55 kDa) is consistent with commercial Blincyto.

The development and validation of a highly sensitive ELISA for GP101 quantitation provides a critical tool for preclinical and clinical studies. We developed a precise and accurate method with a wide range of GP101 concentrations, 15pg/mL to over 500 pg/mL. We found that 25% mouse serum is the optimal matrix for ELISA performance, offering the highest signal-to-noise (S/N) ratios and maintaining reliable precision and accuracy for PK studies where low sensitivity is a target.

The validated ELISA method exhibited excellent sensitivity, with a lower limit of quantitation (LLOQ) of 40 pg/mL in 25% mouse serum. Overall, this ELISA provides a highly sensitive and reproducible method for detecting GP101 in serum. It is likely that a similar approach could be used for the detection of other similar bispecific diabodies.

## Acknowledgments

The authors sincerely thank D. Flasher and C. Murphy-Quinn for their contributions to the experimentation and C. Fraser for assistance in manuscript preparation.

